# HITS-CLIP analysis of human ALKBH8 points towards its role in tRNA and noncoding RNA regulation

**DOI:** 10.1101/2022.01.17.476611

**Authors:** Ivana Cavallin, Marek Bartošovič, Tomas Skalický, Praveenkumar Rengaraj, Martina Christina Schmidt-Dengler, Aleksej Drino, Mark Helm, Štěpánka Vaňáčová

## Abstract

Transfer RNAs acquire a large plethora of chemical modifications. Among those, modifications of the anticodon loop play important roles in translational fidelity and tRNA stability. Four human wobble U containing tRNAs obtain 5-methoxycarbonylmethyluridine (mcm^5^U_34_) or 5-methoxycarbonylmethyl-2-thiouridine (mcm^5^s^2^U_34_), which play a role in decoding. This mark involves a cascade of enzymatic activities. The last step is mediated by Alkylation Repair Homolog 8 (ALKBH8). In this study, we performed a transcriptome-wide analysis of the repertoire of ALKBH8 RNA targets. Using a combination of HITS-CLIP-seq and RIP-seq analyses, we uncover ALKBH8-bound RNAs. It targets an additional wobble U-containing tRNA tRNA^Lys^(UUU). More interestingly, the spectrum of bound RNAs includes other types of non-coding RNAs, such as C/D box snoRNAs, 7SK RNA or some miRNAs, respectively.

## Introduction

Chemical RNA modifications have emerged as an important regulator of gene expression in many living systems. In tRNAs, modifications of the anticodon residues allow the decoding capacity to expand. This includes A to I editing by ADATs, m^5^C mediated by NSUN2 and TRDMT1, queuosinylation, and mcm^5^U (Agris et al. 2018). Wobbling is the pairing of the third codon nucleotide with the first anticodon nucleotide, termed the wobble nucleotide, by a non-Watson-Crick bond. This allows recognition of non-cognate codons (Crick 1966), and consequently enables substitution of a missing tRNA decoder or regulates translation (Grosjean et al. 2010; Endres et al. 2015b). In bacteria and organelles, there are cases of superwobbling where an unmodified wobble uridine can decode all four nucleotides (Rogalski et al. 2008; Inagaki et al. 1995; Alkatib et al. 2012). However, in the eukaryotic cytosol, wobble uridine is almost invariably modified (Schaffrath and Leidel 2017; Yokoyama et al. 1985). Modified uridine is most commonly found in split codon boxes, and it is presumed that it prevents mistranslation of the pyrimidine encoded amino acid (Johansson et al. 2008).

To date, decoding by wobble uridine has been studied mainly in yeast. Analysis of purified tRNAs identified different modifications of wobble uridines (Fig 1A, **S1A**) (Smith et al. 1973; Kobayashi et al. 1974; Kuntzel et al. 1975; Randerath et al. 1979; Yamamoto et al. 1985; Keith et al. 1990; Glasser et al. 1992; Szweykowska-Kulinska et al. 1994; Lu 2005; Johansson et al. 2008). Trm9 in *Saccharomyces cerevisiae* performs the final step of wobble uridine modification, methylation of 5-carboxymethyluridine (cm^5^U) and 5-carboxymethyl-2-thiouridine (cm^5^s^2^U) to 5-methoxycarbonylmethyluridine (mcm^5^U) and 5-methoxycarbonylmethyl-2-thiouridine (mcm^5^s^2^U), respectively (Kalhor and Clarke 2003). The first step in the wobble uridine modification is catalyzed by the Elongator complex. The Elongator is composed of six subunits Elp1 – Elp6 (Huang 2005; Dauden et al. 2017). The Elp3 subunit catalyzes the transfer of the carboxymethyl (cm) residue to the 5-position of uridine (Selvadurai et al. 2014). Trm9 then converts cm^5^ to mcm^5^ (Kalhor and Clarke 2003; Songe-Moller et al. 2010; Fu et al. 2010a). Carbamoylmethyl (ncm^5^) modification has also been identified on wobble uridines, particularly in mutants with defective Trm9 (Johansson et al. 2008; Chen et al. 2011). The enzyme responsible for the conversion to ncm^5^ has not been uncovered and it is unclear whether this modification represents an alternative pathway or a step preceding the conversion to mcm^5^. However, in *Arabidopsis* mutants depleted of AtTRM9, the level of ncm^5^ corresponded to the level in wild type (Leihne et al. 2011). This suggests that at least some pathways to the formation of these modifications might be distinctive for plants. The other two modifications found on wobble uridines together with mcm^5^ are 2-thiolation and 2'-O-methylation. Ncs2/Ncs6 catalyze the 2-thiolation of uridine (Huang et al. 2008; Nakai et al. 2008; Leidel et al. 2009; Noma et al. 2009). Its level declined in the Elongator and Trm9 mutants (Nakai et al. 2008; Noma et al. 2009). Trm7/Trm734 methylate the 2'-O of ribose in tRNA^Leu^(ncm^5^UmAA) (Pintard 2002; Guy et al. 2012). This ribose modification appears to be independent of base modifications, since Um could be found in wild type cells, and tRNA^Leu^(ncm^5^UAA) could be found in Trm7 deletion mutants (Guy et al. 2012).

**FIG. 1.**
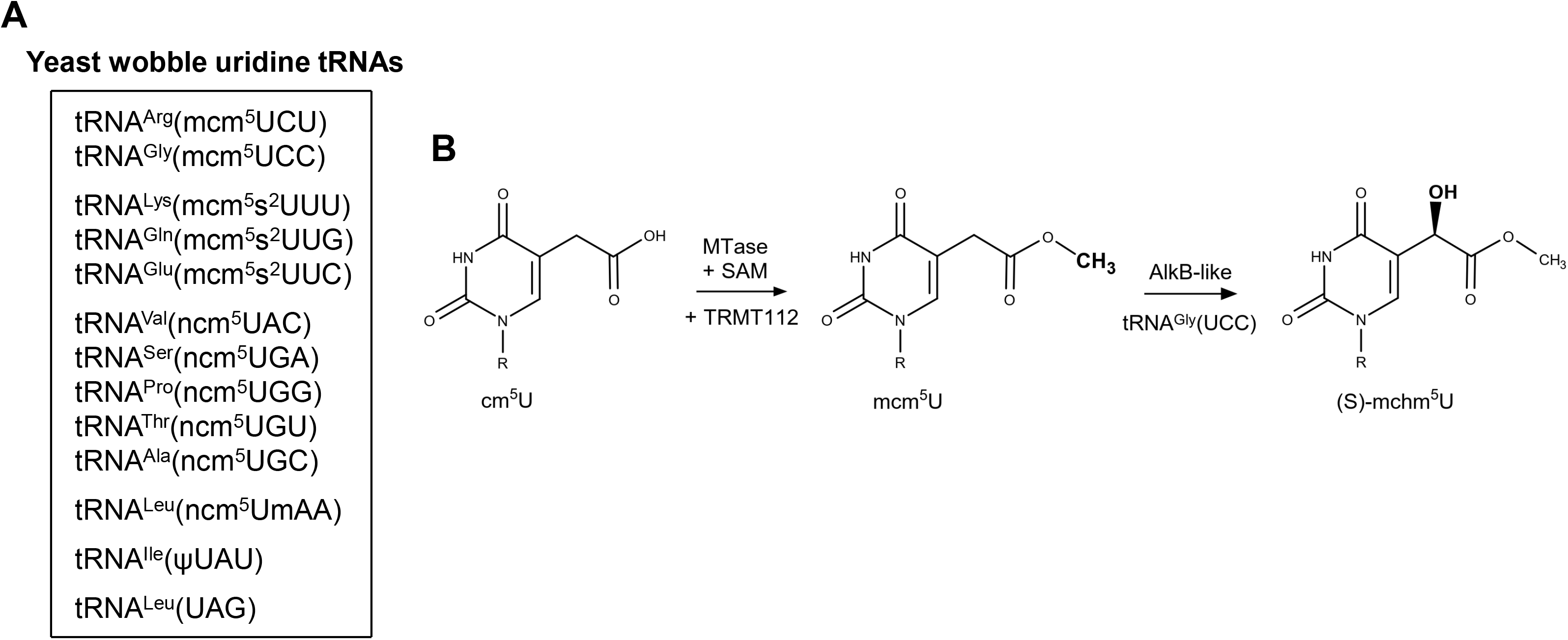
Wobble uridine tRNA modifications and ALKBH8 activities. (A) Known wobble uridine modifications in yeast. (B) The ALKBH8 reaction mechanism. The MTase domain of ALKBH8 together with the cofactor TRMT112 uses S-adenosyl-L-methionine (SAM) to convert cm^5^U into mcm^5^U. In case of tRNA^Gly^(UCC), the AlkB-like domain subsequently performs hydroxylation into (S)-mchm^5^U.

The human AlkB homolog (ABH) family includes nine enzymes – ALKBH1 to ALKBH8 (Wei et al. 1996; Duncan et al. 2002; Aas et al. 2003; Tsujikawa et al. 2007) and FTO (Gerken et al. 2007). Among them, ALKBH8 is the ortholog of Trm9 (Songe-Moller et al. 2010; Fu et al. 2010a). ALKBH8 has a specific domain composition. Besides the central AlkB-like domain that is common for this family, it contains the N-terminal RNA recognition motif (RRM) and the C-terminal methyltransferase (MTase) domain (**Fig S1B**) (Pastore et al. 2012). The MTase domain requires the cofactor TRMT112 to catalyze wobble uridine methylation (Songe-Moller et al. 2010; Fu et al. 2010a).

Increased levels of ALKBH8 were found in high-grade invasive bladder cancer (Shimada et al. 2009; Ohshio et al. 2016), while another study reported downregulation in breast, bladder, colorectal, cervix, and testicular carcinoma (Begley et al. 2013). The impairment of ALKBH8 catalytic function by truncating mutations in the C-terminal region was associated with intellectual disability (Monies et al. 2019). A point mutation in *IKBKAP* that encodes ELP1 causes familial dysautonomia (Anderson et al. 2001; Slaugenhaupt et al. 2001), while other mutations are linked to bronchial asthma (Takeoka et al. 2001). Mutated ELP3, ELP4, and FTSJ1, which is the ortholog of Trm7, were associated with amyotrophic lateral sclerosis, Rolandic epilepsy, and mental retardation, respectively (Simpson et al. 2009; Strug et al. 2009; Freude et al. 2004; Ramser 2004).

Modifications in human tRNAs may differ from yeast modifications. Human tRNA^Arg^(UCU) was found to contain mcm^5^s^2^ in contrast to mcm^5^ in yeast (Songe-Moller et al. 2010), and the stop codon-recoding tRNA^SeC^(UCA) is present in two subgroups of wobble modification: mcm^5^U and mcm^5^Um (Songe-Moller et al. 2010). Furthermore, the AlkB-like domain of ALKBH8 was shown to hydroxylate mcm^5^U to (S)-methoxycarbonylhydroxymethyluridine ((S)-mchm^5^U) in tRNA^Gly^(UCC) (Fig 1B) (Fu et al. 2010b; van den Born et al. 2011). Its diastereomer (R)-mchm^5^U is present in tRNA^Arg^(UCG), but it remains unclear which enzyme mediates this hydroxylation.

Studies of human ALKBH8 tRNA targets have been largely directed by data available from yeast studies. To date, ALKBH8 RNA binding and enzymatic activity have been demonstrated at the following specifically selected tRNAs, tRNA^Arg^(UCU), tRNA^Glu^(UUC), tRNA^SeC^(UCA), and tRNA^Gly^(UCC) (Songe-Moller et al. 2010; Fu et al. 2010a, 2010b; van den Born et al. 2011). Other wobble U-containing tRNAs have not been addressed to date. In this study, we present an unbiased study of human ALKBH8 RNA targets by using high-throughput sequencing of crosslinked and immunoprecipitated RNAs (HITS-CLIP-seq). Our results reveal the previously characterized tRNAs as well as several other tRNAs. Interestingly, the study also suggested binding of ALKBH8 to several other types of RNA.

## Materials and Methods

### Preparation of the FLAG-ALKBH8 expression construct

The synthetic sequence of the 3 x FLAG tag (**Table S1**) was inserted between the KpnI and NotI sites of the vector pcDNA5/FRT/TO (ThermoFisher Scientific) to prepare a DNA construct for N-terminal FLAG fusions. The full-length (1994 bp) *ALKBH8* coding region was amplified using primers ALKBH8_fwd and ALKBH8_rev (**Table S1**). As a template for PCR, we used oligo dT-primed cDNA synthesized from total RNA isolated from HEK293 T-Rex Flp-In cells. The PCR product was inserted between the NotI and XhoI sites of pcDNA5/FRT/TO/FLAG to obtain the N-terminal 3 x FLAG fusion gene. The final construct was verified by sequencing.

### Cell culture and stable cell line preparation

Human Embryonic Kidney (HEK293) T-Rex Flp-In (ThermoFisher Scientific) cells were grown in DMEM supplemented with 10% FBS in an atmosphere of 5% CO2, 37°C. At 70% confluency, cells were cotransfected with 1 μg of pcDNA5/FRT/TO/3xFLAG-ALKBH8 and 10 μg of pOG44 using TurboFect Transfection Reagent (ThermoFisher Scientific). After 24 hours, cells were transferred to a 15-cm dish and recombinants were selected in the presence of 100 μg/ml of hygromycin B. When individual cell colonies were grown, the colonies were tested for doxycyclin-inducible expression of 3 x FLAG-ALKBH8 and zeocin sensitivity.

### Western blot

Proteins were resolved by SDS-PAGE and transferred to a nitrocellulose membrane. Primary antibodies used in this study are: anti-FLAG (1:3000, Sigma Aldrich F3165), anti-tubulin (1:10000, Sigma Aldrich T6074), anti-fibrillarin (1:1000, Abcam ab5821), anti-ALKBH8 (1:500, Abcam ab113512). Washes and incubation of HRP-conjugated secondary antibodies (1:5000, Promega) were performed in PBST (0.05% Tween 20 in PBS). Pierce ECL Western Blot Substrate (ThermoFisher Scientific) was used for visualization.

### Immunofluorescence

Cells grown on polyethyleneimine-coated coverslips were fixed with 3.7% paraformaldehyde for 30 min and permeabilized with 0.2% Triton X-100 in PBS for 20 min. 1 hour of blocking with 5% horse serum in PBS was followed by incubation with anti-FLAG primary antibody (1:500, Sigma Aldrich F3165) in 3% horse serum in PBS for 1 hour. Alexa594 secondary antibody (1:1000, Invitrogen A-21203) incubation was performed together with DAPI (1:500, Sigma) in 3% horse serum in PBS for 30 min. All washes were performed in PBS and the final wash was in PBST. Coverslips were mounted on microscopy slides in Mowiol 4-88 (Sigma) with DABCO. The imaging for the localization overview was performed on the Zeiss AxioImager.Z2-Zen upright fluorescence microscope with the objective Plan-Apochromat 63x/1.40 Oil.

### Subcellular fractionation

Subcellular fractionation was performed as previously published (Ustianenko et al. 2016). Briefly, HEK293 T-Rex Flp-In cells were permeabilized in the dish with digitonin (45 μg/ml digitonin, 10 mM DTT, 10 mM MgCl_2_ in 1 x NEH buffer). The released cytoplasmic fraction was collected and cleared by centrifugation. Cell residues in the dish were scraped in buffer 2 (150 mM NaCl, 1% NP-40 50 mM HEPES pH 7.4). The nuclear fraction was harvested by centrifugation.

### HITS-CLIP-seq protocol

HITS-CLIP was performed in duplicates as previously described in (Bartosovic et al. 2017). Briefly, cells induced with 0.1 mg/ml of doxycycline were UV-crosslinked (150 mJ/cm at 254 nm) and lysed in 100 mM NaCl, 50 mM Tris pH 7.4, 1% NP-40, 0.1% SDS, 0.5% sodium deoxycholate, and 1 x EDTA-free Complete Protease Inhibitor Cocktail (Roche). After brief sonication, the lysates were treated with Turbo DNase (Fermentas) and RNase I (Ambion). Protein G Dynabeads (ThermoFisher Scientific) were coupled with anti-FLAG antibody (Sigma) and used for the immunoprecipitation of FLAG-ALBH8 complexes. The beads were then washed with high salt buffer (lysis buffer with 1 M NaCl and 1 mM EDTA) and PNK buffer (20 mM Tris pH 7.4, 10 mM MgCl_2_, 0.2% Tween 20). RNA was treated with Fast AP Thermosensitive Alkaline Phosphatase (Fermentas), and the L3 DNA linker (**Table S1**) was ligated to RNA by T4 RNA Ligase 2 (NEB). The beads were washed with high salt buffer and PNK buffer. RNA was labeled with gamma-^32^P-ATP using T4 PNK (NEB). After washing with PNK buffer, the elution from the beads was performed by incubation at 95°C for 5 min. The eluted protein-RNA complexes were resolved on a 4-12% NuPage gel (ThermoFisher Scientific), transferred to a nitrocellulose membrane (BioRad) and the area corresponding to the migration of ALKBH8 and 20 kDa above was cut from the membrane. The membrane was cut into pieces and PK buffer (100 mM Tris pH 7.4, 50 mM NaCl, 10 mM EDTA) was added together with Proteinase K (NEB). After 90 min of incubation at 37°C, an equal volume of PK buffer with 7 M urea was added and RNA was extracted by incubation at 55°C for one hour. RNA was purified by phenol/chloroform extraction and ethanol precipitation. The RNA linker L5 (**Table S1**) was ligated with T4 RNA ligase (NEB). RNA was purified by phenol/chloroform extraction. SuperScript III (ThermoFisher Scientific) and random hexamers were used for reverse transcription and PCR amplification was carried out using 25 cycles and GoTaq polymerase (Promega) with linker specific primers. The quality of the PCR libraries was analyzed by Bioanalyzer, and high-throughput sequencing was performed in the 75bp single-end mode on Illumina HiSeq 2000 at the Vienna BioCenter (VBCF) Sequencing facility.

### RNA immunoprecipitation and preparation of cDNA library (RIP-seq)

FLAG-ALKBH8 expression was induced with 0.1 mg/ml doxycycline at 70% confluency in two 10 cm plates. 18 hours later, cells were harvested, washed twice with 1 x PBS, and lysed in 1.4 ml of lysis buffer (50 mM Tris pH 8.0, 150 mM NaCl, 1% Triton X-100, 1 mM DTT, 1 x EDTA-free Complete Protease Inhibitor Cocktail (Roche) and 110 U/ml of RNasin (Promega)) rolling for 30 min at 5°C. The lysate was treated with 2 U/ml Turbo DNase (Fermentas) for 15 min at 37°C. The insoluble fraction was removed by centrifugation at 15000 g for 15 min, the supernatant was incubated with 1/14 volume of free Protein G Dynabeads (ThermoFisher Scientific) rolling for 1 hour at 5°C to remove sticky proteins and RNAs. The precleared lysate was then transferred to Protein G Dynabeads coupled with anti-FLAG antibody (Sigma) and the mixture was incubated for 2 hours at 5°C on a roller. The beads with bound proteins were washed three times with 1 ml of lysis buffer. RNA was isolated using TriPure Isolation Reagent (Roche) according to the manufacturer’s protocol, precipitated in an equal volume of isopropanol and 7.5 μg/ml of GlycoBlue (Ambion) at −20°C, washed with 1 ml of 85% ethanol and the pellets were resuspended in 10 μl H_2_O.

The L3 DNA linker at 0.5 μM concentration (**Table S1**) was ligated to RNA in a 20 μl reaction using 5 U/μl of RNA Ligase Truncated K227Q (NEB) in the presence of 15% PEG8000 and 1 U/μl RNasin (Promega) at 18°C overnight and then at room temperature for 1 hour and inactivated at 65°C for 20 min. RNA was purified by phenol/chloroform extraction and ethanol precipitation and the pellets were resuspended in 6 μl H_2_O. The L5 RNA linker at 1 μM concentration (**Table S1**) was ligated to RNA in a 20 μl reaction using by 1 U/ μl of RNA Ligase 1 (NEB) in the presence of 15% PEG8000, 1 U/μl RNasin (Promega) and 1 mM ATP at 16°C overnight. RNA was purified by phenol/chloroform extraction and ethanol precipitation. SuperScript IV (Invitrogen) was used for reverse transcription and the resulting cDNA was treated with 1.25 U/μl RNase H (Invitrogen). cDNAs were amplified by 15 PCR using GoTaq G2 polymerase (Promega) and TruSeq_adaptor_fwd and TruSeq_adaptor_rev primers (**Table S1**). The PCR products were resolved on a 2% agarose gel and the fragments migrating within the range of 200 to 330 nt were excised and the DNA was extracted using the Gel Extraction Kit (Qiagen) according to the manufacturer’s instructions. 125-bp pair-end sequencing was performed on Illumina HiSeq V4 PE125 at the Vienna BioCenter Core Facilities.

### Northern blot analysis

RNA was separated on a 16% polyacrylamide gel and transferred to the Amersham Hybond-N+ membrane (GE Healthcare). Probes tRNA-Glu-TTC, tRNA-Lys-TTT, tRNA-Lys-CTT, and tRNA-Gly-CCC (**Table S1**) were radioactively labeled with gamma-^32^P-ATP using T4 PNK (NEB). Pre-hybridization and hybridization were performed in OligoHyb Buffer (Applied Biosystems) at 50°C. The membrane was washed twice with 300 mM NaCl, 30 mM tri-sodium citrate, 0.1% SDS, pH 7.0, and twice with 15 mM NaCl, 1.5 mM tri-sodium citrate, 0.1% SDS, pH 7.0. The radioactive signal was detected by autoradiography.

### Bioinformatics analyses of high-throughput sequencing data

Sequencing quality control was performed with the FastQC tool (Andrews 2010) and preprocessing, including the removal of sequencing adapters, and the removal of low-quality reads (Q<25) and bases was achieved with Trimmomatic (Bolger et al. 2014). High quality reads were mapped to a high confidence tRNA dataset obtained from GtRNAdb using the Smith-Waterman algorithm for optimal local alignment in R (ver. 3.6.3) and Biostrings package (ver. 2.52.0) with the following settings: pairwiseAlignment (target_seq, query_seq, type = “local”, substitutionMatrix = nucleotideSubstitutionMatrix (match = 1, mismatch = −1), gapOpening = 10, gapExtension = 0.1). Only uniquely mapped reads that aligned at least in 66 % of their length and had a maximum of 3 mismatches were used for later analyses.

Reads that did not map uniquely to this dataset were mapped to the human genome version GRCh38.p12 by STAR (Dobin et al. 2013), a splice-aware mapping program using ENSEMBL recommended settings, with maximum intron size = 10 kb, 3 nt or 0.03% mismatches and allowed soft clipping. Uniquely mapped reads were identified using Samtools (Li et al. 2009), and CLIP reads were categorized and annotated according to the priority table (1. rRNA, 2. tRNA, 3. snoRNA, 4. snRNA, 5. miRNA, 6. mRNA exon, 7. mRNA intron, 8. lincRNA, 9. repetitive elements 10. antisense mRNA, 11. unannotated). In the next step, the genome with mapped reads was dissected into 10 bp windows and the coverage for each CLIP library was calculated using the bamcoverage utility from the Deeptools2 package (Ramírez et al. 2016). Data from both mapping to GtRNAdb and the human genome were normalized using the counts per million (CPM) method. Identification of the crosslinking site in HITS-CLIP data mapped to the human genome was performed with Piranha (Uren et al. 2012). To identify the highest probable targets of ALKBH8 in addition to tRNA, we have combined the results of annotation, coverage calculation, and peak calling using the intersectBed tool from the BEDtools suite (Quinlan and Hall 2010) and sorted them according to CPM and identified peaks. The heatmap showing the binding regions on selected tRNA target genes was generated using the Deeptools2 package with the scaled region set to 74 nucleotides. To identify the non-templated 3' CCA tails from RIP-seq data, the reads exploited were the first ones of the paired reads that mapped uniquely into genomic regions of selected tRNAs. The soft-clipped parts were extracted. The CPM normalized numbers of non-templated 3' tails and of sequences without any extension were counted using a custom Python script. We considered the sequences as containing non-templated 3' tails only when they were ending with C, CC, CA, CCC, CCA, CCAA, or CCACCA.

### Purification of cellular tRNA^Lys^(UUU) and tRNA^Glu^(CUC)

tRNA^Lys^(UUU) and tRNA^Glu^(CUC) were purified according to (Drino et al. 2020). Briefly, total RNA was isolated from three 15 cm plates of HEK293T cells grown to 90% confluency using Trizol and isopropanol precipitated. Small RNA fractions, containing tRNAs, were eluted in 40-60% of IEX buffer B (20 mM Tris, 1 M NaCl, 10 mM KCl, 1.5 mM MgCl_2,_). For ion exchange chromatography, a HiTrap Q FF anion exchange chromatography column (1 ml, GE Healthcare) was used on an ÄKTA-FPLC (GE Healthcare) at 4°C. Eluted fractions between 400-600 mM NaCl were collected and immediately precipitated using isopropanol at −20°C. Eluted small RNA fractions were pooled and re-suspended in water, supplemented with 10 mM MgCl_2_ and renatured by incubation at 75°C for three minutes, immediately cooled on ice and mixed with 20 ml of binding buffer (30 mM HEPES KOH, 1.2 M NaCl, 10 mM MgCl_2_). For RNA affinity capture, 80 µg of a 5' amino-modified DNA oligonucleotide (**Table S1**) complementary to target tRNA (IDT) was covalently coupled to a HiTrapTM NHS-activated HP column (1 ml, GE Healthcare). Binding to the NHS column was performed by recirculation of the small RNA fraction over the complementary NHS columun for 3 hours at 65°C. The columns were washed with wash buffer (2.5 mM HEPES KOH, 0.1 M NaCl, 10 mM MgCl_2_) and tRNAs were eluted by submerging the column in a water bath at 75°C in elution buffer (0.5 mM HEPES-KOH, 1 mM EDTA), precipitated with isopropanol at −20°C and the pellets were resuspended in water. Purified tRNAs were further gel-purified (8% urea-PAGE in 0.5 x TBE) using RNA gel extraction buffer (0.3 M NaOAc, pH 5.2, 0.1% (v/v) SDS, 1 mM EDTA) and immediately precipitated with isopropanol and resuspended in water.

### LC-MS/MS analysis of tRNA modifications

LC-MS analysis was performed as described previously (Thüring et al. 2016). 500 ng of RNA was digested to the nucleoside level using 0.6 U nuclease P1 from *P. citrinum* (Sigma-Aldrich), 0.2 U snake venom phosphodiesterase from *C. adamanteus* (Worthington), 0.2 U bovine intestine phosphatase (Sigma-Aldrich), 10 U benzonase (Sigma-Aldrich) and 200 ng Pentostatin (Sigma-Aldrich) in 5 mM Tris (pH 8) and 1 mM magnesium chloride for two hours at 37°C. 450 ng of digested RNA was spiked with 50 ng of internal standard (^13^C stable isotope-labeled nucleosides from *S. cerevisiae*) and analyzed by LC-MS (Agilent 1260 Infinity system in combination with an Agilent 6470 Triple Quadrupole mass spectrometer equipped with an electrospray ion source (ESI)). For HPLC separations, a C18 reverse HPLC column (Synergi^TM^ 4 µM particle size, 80 Å pore size, 250 × 2.0 mm; Phenomenex) was used. A gradient solvent system consisting of solvent A (5 mM ammonium acetate buffer, pH 5.3, adjusted with acetic acid) and solvent B (LC-MS grade acetonitrile, Honeywell) was used and the compounds were eluted with a linear gradient of 0-8% solvent B over 10 min, followed by 8-40% solvent B over 10 min. Initial conditions were regenerated with 100% solvent A for 10 min. The flow rate was 0.35 ml/min and the separations were performed with the column temperature maintained at 35°C. The four main nucleosides were detected photometrically at 254 nm using a diode array detector (DAD). Electrospray ionization mass spectra were recorded in positive polarity at a gas temperature of 300°C, gas flow of 7 L/min, nebulizer pressure of 60 psi, sheath gas temperature of 400°C, sheath gas flow of 12 L/min and capillary voltage of 3000 V. Agilent MassHunter software was used in the dMRM (dynamic multiple reaction monitoring) mode.

## Results and Discussion

### Identification of ALKBH8 targets by HITS-CLIP-seq

To identify ALKBH8 RNA targets, we aimed to perform the HITS-CLIP-seq analysis (Ule et al. 2005; König et al. 2010; Bartosovic et al. 2017). We first prepared a stable cell line of HEK293 T-Rex Flp-In (293T) with inducible expression of N-terminally tagged ALKBH8 (FLAG-ALKBH8) (Fig 2A). Both, the FLAG-ALKBH8 and endogenous ALKBH8 exhibited cytoplasmic localization (Fig 2B, 2C), which is in agreement with the previously reported localization of episomally expressed GFP-ALKBH8 (Fu et al. 2010a; Pastore et al. 2012). FLAG-ALKBH8 formed strong protein-RNA complexes after UV irradiation (Fig 2D). We performed three independent experiments. The bioinformatic analysis of HITS-CLIP-seq resulted in 15 798 568 uniquely mapped reads to the human genome (see methods for details). As expected, annotation of mapped reads revealed tRNAs as the most prevalent target (87% of reads), followed by mRNA exons (7%), mRNA introns (3%), repetitive elements (1%), snoRNAs (0.2%) and snRNAs, miRNA and rRNA (all less than 0.1%) (Fig 2B, Supplementary Table2).

**FIG. 2.**
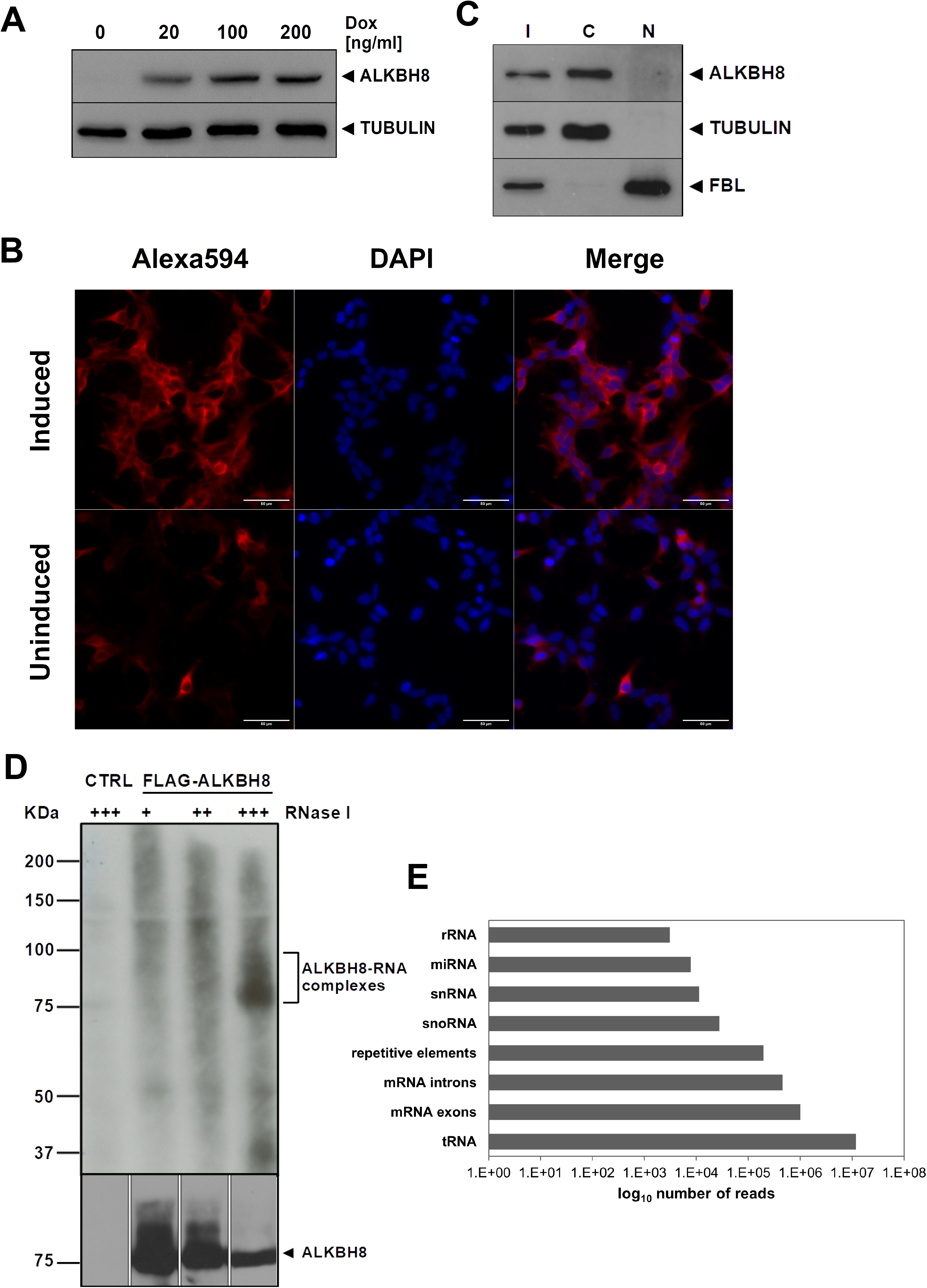
HITS-CLIPseq analysis of ALKBH8 RNA targets in HEK293 T-Rex Flp-In cells identifies a broader RNA binding repertoir. (A) Western blot analysis of the doxycycline-inducible expression of stably integrated 3 x FLAG-ALKBH8. Tubulin was used as a loading control. (B) FLAG-ALKBH8 localizes in the cytoplasm in HEK293T. Immunofluorescence of FLAG-ALKBH8 whose expression was induced by doxycycline and visualized by anti-FLAG and Alexa594 antibodies (red). DNA was stained with DAPI (blue). The scale bar represents 50 μm. (C) ALKBH8 is present mainly in the cytoplasm. Western blot analysis of the total cell lysate (I), cytoplasmic (C), and nuclear (N) fractions of HEK293T cells. ALKBH8 was detected with a specific antibody. Tubulin was used as a cytoplasmic marker and fibrillarin (FBL) as a nuclear marker. (D) ALKBH8-RNA complexes formed by UV treatment. Cellular lysates were treated with an increasing concentration of RNaseI (marked above the gel). CTRL was a negative control of HEK293T in which FLAG-ALKB8 was not expressed. ALKBH8 was immunoprecipitated with anti-FLAG antibody. RNA was 5‘-terminally labeled with δ^32^P-ATP, and complexes were resolved on polyacrylamide and SDS-PAGE denaturing gels, respectively. The upper panel displays the radioactive signal of the RNA detected by autoradiography, and the lower panel is a western blot analysis with anti-FLAG antibody. (E) RNA classes identified by ALKBH8 HITS-CLIP-seq experiments. The graph shows numbers of uniquely mapped reads (x scale).

### ALKBH8 targets a wider repertoire of U_34_-containing tRNAs

The list of previously reported and validated mammalian ALKBH8 substrates included tRNA^Arg^(UCU), tRNA^Glu^(UUC), tRNA^Gly^(UCC), and tRNA^SeC^(UCA) (Fig 3A, top left) (Fu et al. 2010a, 2010b; Songe-Moller et al. 2010; van den Born et al. 2011). Our analysis identified additional tRNAs containing U at position 34 that were not previously reported as ALKBH8 targets in mammals (Fig 3A, top right, **Table S2**). Among those, tRNA^Lys^(UUU) had the highest coverage. tRNA^Lys^(UUU) is modified by the ALKBH8 ortholog Trm9 in yeast where it mediates mcm^5^s^2^U_34_ (Johansson et al. 2008). In their earlier work, Fu et al. did not detect tRNA^Lys^(UUU) among ALKBH8 bound tRNAs (Fu et al. 2010a). Other new potential tRNAs harboring wobble U targets are tRNA^Gln^(UUG), tRNA^Leu^(UAA), tRNA^Leu^(UAG), and tRNA^Val^(UAC). In yeast, tRNA^Gln^(UUG) contains mcm^5^, while tRNA^Leu^(UAA) and tRNA^Val^(UAC) contain ncm^5^ modification for which the responsible enzymatic activities have not yet been identified (Randerath et al. 1979; Kalhor and Clarke 2003; Johansson et al. 2008). No wobble U modification has been reported for yeast tRNA^Leu^(UAG).

**FIG. 3.**
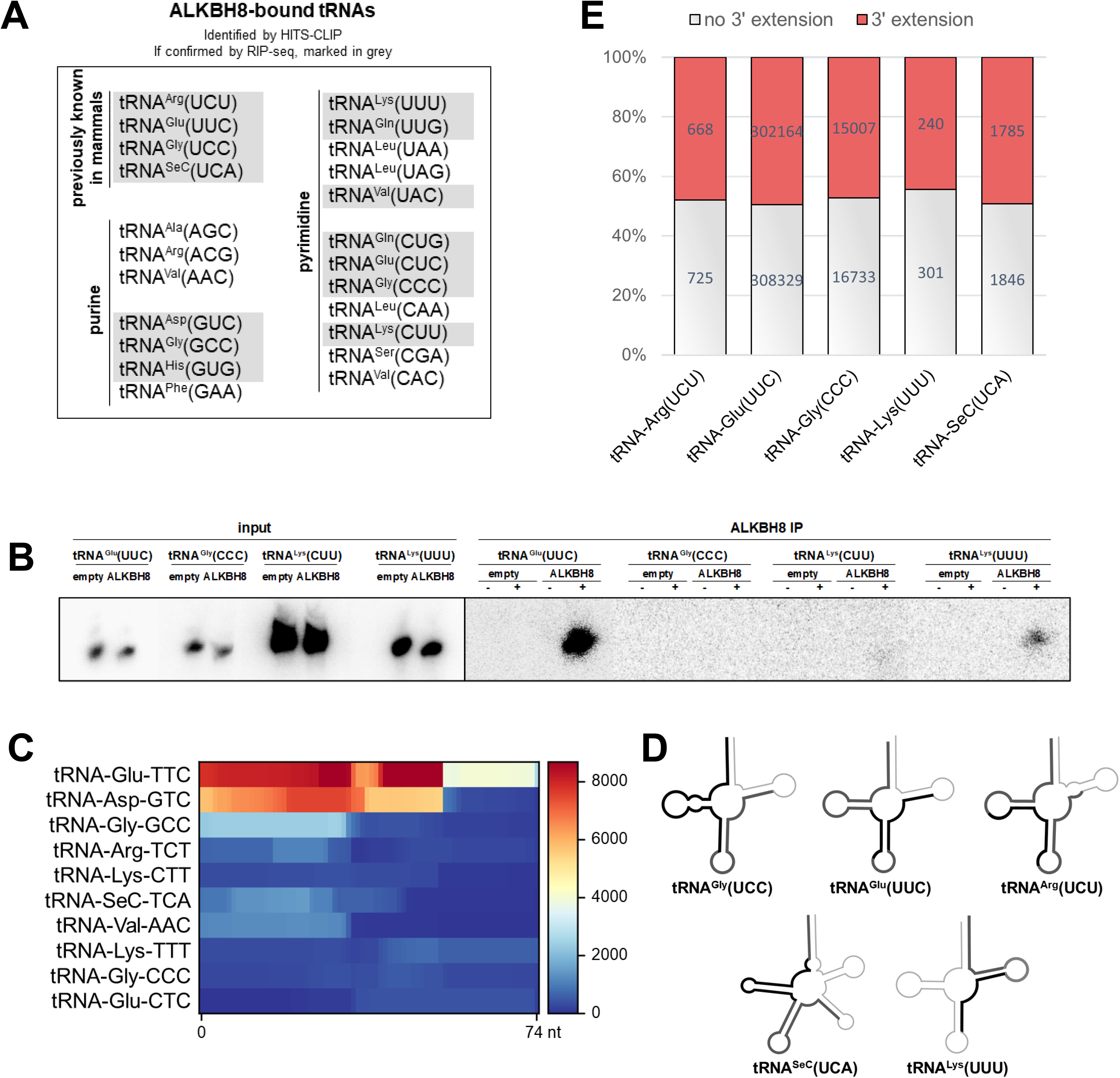
Analysis of ALKBH8-bound tRNAs identified by HITS-CLIP. (A) List of tRNAs identified by HITS-CLIP-seq. The displayed tRNAs are the top-scoring tRNAs according to the CPM in **Table S2**. The tRNAs also observed by ALKBH8 RIP-seq analysis are highlighted in gray. The first four (top left) are the previously known ALKBH8 tRNA targets. tRNAs are displayed according to the identity of the wobble nucleotide. (B) Northern blot analysis of selected tRNAs co-precipitated with ALKBH8. Empty are HEK293T cells without the integrated FLAG-ALKBH8 gene. ALKBH8 are stable cell lines that express FLAG-ALKBH8. Input is RNA isolated from whole cell extracts. ALKBH8 IP are RNAs co-precipitated with affinity purified ALKBH8. Beads lacking anti-FLAG antibody were used as a control for non-specific binding of tRNA (-). FLAG-coupled beads are marked as +. DNA oligo probes for individual tRNAs were 5 terminal labeled by γ^32^P-ATP. The RNAs were resolved on 20% polyacrylamide gel and the radioactive signals detected by autoradiography. (C) Heatmap representation of ALKBH8 binding to tRNAs as revealed by the HITS-CLIP analysis. The strong signal of tRNA-Gly-TCC was omitted in order to decrease the scale of the heatplot. All gene lengths were scaled to an equal length of 74 nt on the X axis. The color scale on the right represents the CPM normalized number of mapped reads to a region. (D) Schematic representation of ALKBH8 binding to the secondary structure of tRNA. The areas with the highest mapping are highlighted by a thick black line, and the lower number of mapped reads is marked in thick grey. (E) Proportions of CPM normalized values of non-templated 3-terminal extensions in most represented tRNAs detected by RIP-seq. The red bars include all the different versions of 3 extensions (C, CC, CCA, see details in Figure S1D). The number of tRNA reads without any non-templated 3-terminal extensions are in gray.

To further validate ALKBH8 binding to tRNA^Lys^(UUU), we analyzed RNAs co-precipitating with FLAG-ALKBH8 (RIP) without stabilization of the binding by crosslinking (**Fig S1C**). Tagged ALKBH8 was immunopurified with FLAG antibody and co-precipitated RNAs were analyzed by northern blot. The probe for tRNA^Glu^(UUC) was used as a positive control of an established target (Fu et al. 2010a). Two tRNAs served as negative controls: tRNA^Lys^(CUU) that appeared among CLIP targets (Fig 3A) but was previously reported as a non-target (Fu et al. 2010a), and tRNA^Gly^(CCC) that also appeared among CLIP targets and does not contain any uridines in the anticodon. Northern blot analysis revealed a strong signal for tRNA^Glu^(UUC) and a weaker signal for tRNA^Lys^(UUU) in the ALKBH8 RIP eluate (Fig 3B). No signal was detected for tRNAs with C at the wobble codon position (Fig 3B). This agrees with the results of the HITS-CLIP-seq, where tRNA^Glu^(UUC) was among the most prominent targets, while tRNA^Lys^(UUU) had ten times lower coverage (**TableS2**).

To verify the HITS-CLIP-seq results on a larger scale, we performed an ALKBH8 RNA immunoprecipitation and sequencing (RIP-seq) analysis. The advantage of this experiment is the absence of RNaseI treatment, which retains information about the full length of the bound small RNAs. We obtained 5 063 455 uniquely mapped reads under strict mapping conditions (1 mismatch allowed). The annotation of the mapped reads revealed a high overlap with the HITS-CLIP-seq data. The most enriched co-precipitated tRNAs included tRNA^Arg^(UCU), tRNA^Glu^(UUC), tRNA^Gly^(UCC), tRNA^SeC^(UCA) and tRNA^Lys^(UUU) (Fig 3A, **Table S2**). Other U_34_-containing tRNAs identified were tRNA^Gln^(UGC), tRNA^Val^(UAC), tRNA^Ile^(UAU) and tRNA^Arg^(UCG) (**Table S2**).

The HITS-CLIP-seq analysis also identified tRNAs with residues other than U at position 34. These include tRNA^Ala^(AGC), tRNA^Arg^(ACG), tRNA^Val^(AAC), tRNA^Asp^(GUC), tRNA^Gly^(GCC), tRNA^His^(GUG), tRNA^Phe^(GAA), tRNA^Gln^(CUG), tRNA^Glu^(CUC), tRNA^Gly^(CCC), tRNA^Leu^(CAA), tRNA^Lys^(CUU), tRNA^Ser^(CGA), and tRNA^Val^(CAC) (Fig 3A, **Table S2**). Some of them were also present in the RIP-seq dataset: tRNA^Asp^(GUC), tRNA^Gly^(GCC), tRNA^His^(GUG), tRNA^Gln^(CUG), tRNA^Glu^(CUC), tRNA^Gly^(CCC), and tRNA^Lys^(CUU).

To further test whether these tRNAs are enzymatically modified by ALKBH8, we purified tRNA^Lys^(UUU) and tRNA^Glu^(CUC) from HEK293T and subjected them to LC-MS analysis. The analysis revealed the presence of mcm^5^U and mcm^5^s^2^U in tRNA^Lys^(UUU) (**Fig S2**). On the contrary, these modifications were not detected in tRNA^Glu^(CUC).

The HITS-CLIP-seq analysis revealed a larger spectrum of tRNAs bound to ALKBH8, including tRNAs with nucleotides other than U at the wobble position. Such binding was also uncovered by RIP-seq, although in this case the tRNAs without wobble uridine were significantly less represented than the wobble uridine-containing tRNAs (**Tab S2**). Noticeably, *tRNA-Glu-CUC-1-1* was detected with high counts per million (CPM) also by non-crosslinked RIP. However, the LC-MS analysis did not detect mcm^5^U or mcm^5^s^2^U, marks resulting from ALKBH8 activity. This tRNA manifests up to 97% identity with tRNA^Glu^(UUC), which is a difference of only 2 nt. One of these two nucleotides is located in the D arm, and the other nucleotide is in the anticodon. An error in mapping to tRNA^Glu^(CUC) was excluded by allowing the maximum mismatch of one nucleotide in the whole read and by using only uniquely mapped reads. Therefore, it is possible that ALKBH8 binds both tRNA^Glu^(UUC) and tRNA^Glu^(CUC) with high affinity *in vivo* due to their high sequence identity. The activity of ALKBH8 would then depend on the prior formation of the cm^5^U modification (Kalhor and Clarke 2003). However, it is also possible that binding of ALKBH8 to other tRNAs may have additional functional consequences.

### ALKBH8 binds to mature tRNAs

tRNAs are produced as precursors with 5 - and 3 - trailers that are endonucleolytically cleaved by RNase P and tRNase Z (Guerrier-Takada et al. 1983; Schiffer et al. 2002) and 3 termini are extended with CCA tails that the CCA nucleotidyl transferase adds (Rossmanith et al. 1995). Upon export to the cytoplasm, tRNA is charged with the corresponding aminoacid (Hampel and Enger 1973). Individual internal chemical modifications occur at different steps of tRNA processing. For instance, wybutosine formation at position 37 in yeast is catalyzed after retrograde import to the nucleus after nuclear intron removal in the cytoplasm (Yoshihisa et al. 2007; Ohira and Suzuki 2011). The timing of ALKBH8 activity and possible coordination with other processes is not yet fully understood. Furthermore, only limited information is available on the molecular principles of ALKBH8 RNA specificity and binding, as only the structure of the human RRM domain and the AlkB-like domain in complex with an oligonucleotide (Pastore et al. 2012). Therefore, we analyzed the properties of ALKBH8 binding to tRNAs in greater detail, as revealed by the HITS-CLIP-seq. We plotted the CLIP coverage of highly covered tRNA targets on their genomic loci to display the sites of interaction with ALKBH8. Both wobble U and non-wobble U tRNAs exhibited a predominant signal over the 5-half of the tRNA sequences (Fig 3C, 3D). tRNA^Gly^(UCC) was excluded from Fig 3C due to its strong coverage that masked the read distribution of less covered tRNAs. The reads for the known targets tRNA^Gly^(UCC), tRNA^Glu^(UUC) and tRNA^Arg^(UCU) mapping were extended over the anticodon loop toward the T-arm. The two tRNAs with predominant binding in the tRNA 3 half were tRNA^Lys^(UUU) and tRNA^Glu^(CUC). The genes of tRNA^Arg^(UCU) tRNA contain a 14-nt intron located downstream of position 37 that formed a signal gap in the read mapping to the genomic locus. Even though we are limited to only one example of an intron-containing tRNA, this finding suggested that ALKBH8 targets mature, fully spliced tRNAs.

Since the HITS-CLIP protocol involves RNA cleavage, we investigated the RIP-seq results that preserved the bound full-length tRNA, to inspect the tRNA termini. In particular, we analyzed the status of the 3-terminal non-templated addition of CCA. 50% of the reads that mapped to the known tRNA targets possessed the non-templated nucleotide addition (Fig 3E). These additions were generally composed of one or two non-templated cytidines (**Fig S1D**). 22% of the tRNA reads contained the mature CCA 3-terminus. tRNA^SeC^(UCA) showed a different distribution of 3-additions, where the CCA trinucleotide formed 70% of all additions (**Fig S1D**). However, the overall proportion of reads lacking any non-templated 3-terminus is in contrast to the consensus that tRNA nuclear export occurs only after the CCA addition. The relative affinity of tRNA for exportin-t/RangGTP is 10 times lower when tRNA is without 3-CCA, and 6 times weaker when tRNA carries only 3-CC (Lipowsky et al. 1999). Therefore, ALKBH8 might bind to tRNAs lacking 3-CCA due to the activity of ANKZF1 and ELAC1 (Yip et al. 2019, 2020). The incomplete CCA termini are repaired by a cytosolic population of CCA-transferase (Wolfe et al. 1996). However, we cannot rule out the possibility that the CCA extensions might have been lost during the RIP protocol due to 3 to 5 exoribonucleolytic activities in the cell lysate.

### ALKBH8 binds to other types of RNA *in vivo*

The highest scoring hits in the HITS-CLIP-seq datasets also included other types of RNA (**Table S2**). We identified a number of C/D box snoRNAs, e.g. *SNORD13, SNORD16*, *SNORD18A, SNORD18C*, *SNORD26*, *SNORD34*, *SNORD42B, SNORD43*, *SNORD58C, SNORD72, SNORD83A* and *SNORD83B* (Fig 4A, **Table S2**). In addition to snoRNAs, binding to other ncRNAs was found, such as *7SK RNA*, *RNY5*, the RNA component of vault particles *VTRNA1-2*, *miR186,* and *miR374A*. (Fig 4A, **Table S2**). Many reads were mapped to exonic and intronic regions of protein-coding genes even under highly strict mapping conditions (no or only one mismatch). In all cases, these mRNAs contained a single well-defined peak of 20 to 60 nt in length, which is narrower than the peaks in other classes of RNA (**Table S2**). Further studies will tackle the question of whether ALKBH8 binding to these regions has any functional significance.

**FIG. 4.**
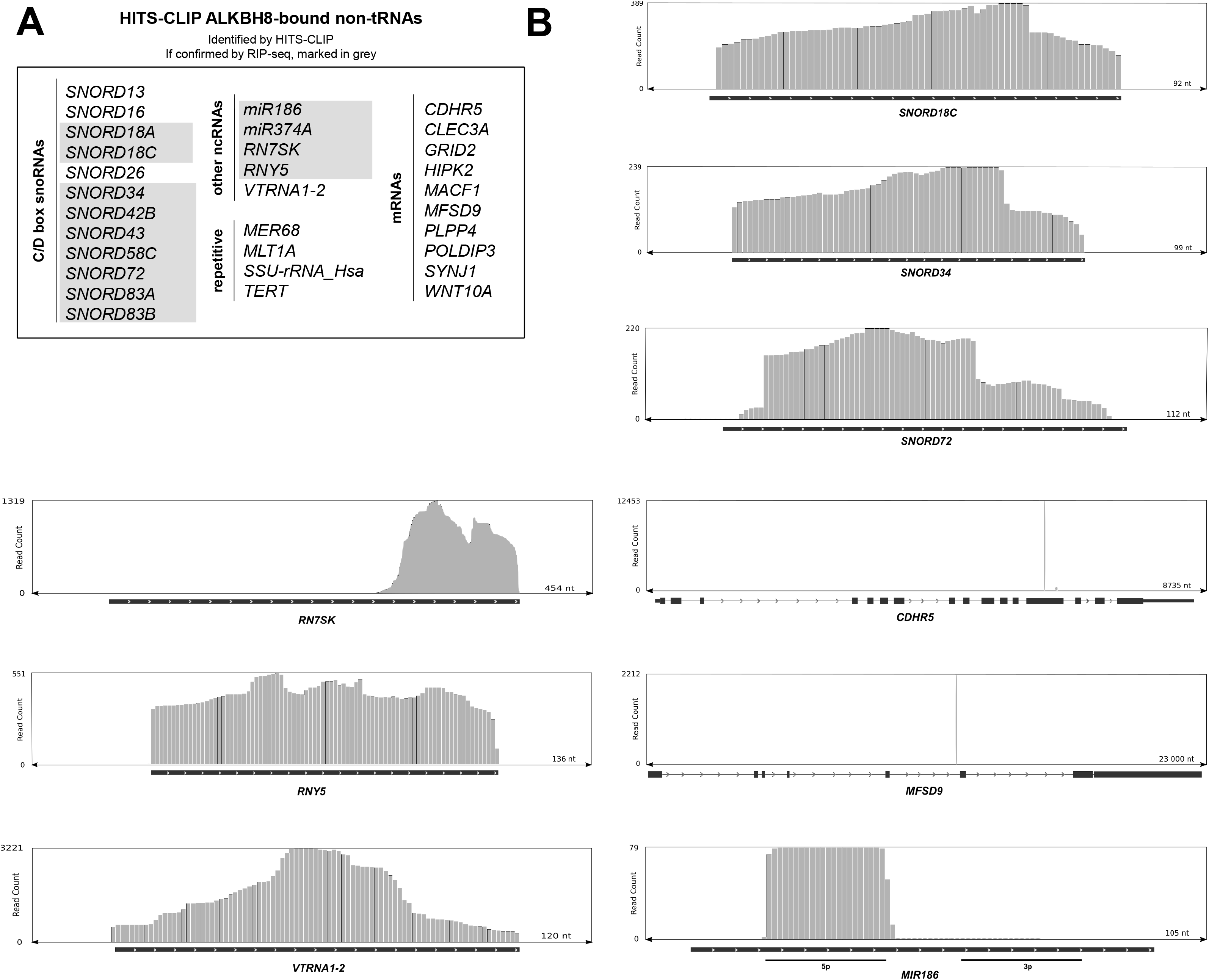
HITS-CLIP-seq analysis of ALKBH8 revealed non-tRNA types of RNA. (A) Top-scoring bound RNAs according to CPM (**Table S2**). Targets that were recapitulated by RIP-seq analysis are colored gray. Genes are listed in numerical and alphabetical order. (B) Representation of ALKBH8 binding to selected genomic regions of RNA.

The RIP-seq dataset included a high number of C/D box snoRNAs (**Table S2**). Comparison of the results of HITS-CLIP-seq and RIP-seq revealed an overlapping set of intron-encoded C/D box snoRNAs (*SNORD18A, SNORD18C*, *SNORD34*, *SNORD42B, SNORD43*, *SNORD58C, SNORD72, SNORD83A,* and *SNORD83B*) and other ncRNAs (*miR186, miR374A*, *7SK RNA*, *RNY5)* (Fig 4A, in gray). Another set of C/D box snoRNAs showed high CPM only in the RIP dataset, e. g. *SNORD63, SNORD101, SNORD117, SNORD118* (**Table S2**). ALKBH8 is localized in the cytoplasm (Fig 1C, E) as other anticodon loop-modifying enzymes such as FTSJ1, the human ortholog of 2-O-methylase Trm7 (Li et al. 2020), or the U_31_-specific pseudouridylase Pus6 (Ansmant et al. 2001). The Elongator, whose activity precedes ALKBH8 on wobble uridines, localizes principally in the cytoplasm and partially in the nucleus (Pokholok et al. 2002; Kim et al. 2002). Despite the location of ALKBH8, there is a striking representation of nuclear C/D box snoRNAs among the identified targets. However, their prominent nucleolar location is not exclusive. *U3 snoRNA* is known to shuttle between the nucleus and the cytoplasm (Leary et al. 2004) and has been reported to be processed into functional miRNAs (Lemus-Diaz et al. 2020). Other snoRNAs were observed in the cytoplasm after oxidative stress (Holley et al. 2015). ALKBH8 is involved in the response to oxidative stress by enhancing the translation of detoxification enzymes containing selenocysteine by catalyzing the modification of tRNA^Sec^(UCA) (Endres et al. 2015a). The partial cytoplasmic localization of snoRNAs in *Arabidopsis thaliana* under fyziological conditions (Streit et al. 2020) indicates a possible cytoplasmic distribution also in other organisms.

Another possible link to ALKBH8 binding to snoRNAs may be the finding that some snoRNAs show activity on tRNAs. For example, human *SNORD97* together with *SCARNA97* catalyzes 2-O-methylation of C_34_ of tRNA^Met^(CAU) (Vitali and Kiss 2019). PAR-CLIP performed on the FBL, NOP56, NOP58 and DKC1 snoRNP proteins showed a small fraction of bound tRNAs (Kishore et al. 2013). It is possible that ALKBH8 and snoRNAs simultaneously interact with tRNA, leading to binding signals in the HITS-CLIP data.

Wobble uridine-like modifications that are known to be catalyzed by ALKBH8 have not been found in other RNA species (Cantara et al. 2011; Boccaletto et al. 2018). However, to our knowledge, mcm^5^U/mchm^5^U has not been examined in detail in a wide range of RNAs. There are enzymes known to have dual specificity on distinct RNA species. For instance, NSUN2 introduces 5-methylcytosine to tRNA at position C_34_ (Brzezicha et al. 2006; Auxilien et al. 2012) and to vault RNAs (Hussain et al. 2013). Furthermore, fluorescence anisotropy assays showed unspecific binding of the ALKBH8/TRMT112 complex to a 17mer composed of random nucleotides with the dissociation constant 350 ± 20 nM (Pastore et al. 2012). This value represents an affinity similar to binding a 17-nt stem-loop derived from a tRNA^Gly^ target that had the dissociation constant of 240 ± 29 nM or to an ALKBH8 aptamer that demonstrated a dissociation constant of 240 ± 50 nM. This supports the hypothesis that ALKBH8 binds and possibly modifies other substrates than tRNA *in vivo*.

Other ncRNAs, namely vault RNA, Y RNA and *7SK RNA*, that were identified in our HITS-CLIP data, had previously been identified in other CLIP-seq experiments with the RNA modifier enzymes Pus7 pseudouridinylase (Guzzi et al. 2018), METTL16 N^6^-adenosine methyltransferase (Warda et al. 2017) or Nsun2 5-methylcytosine transferase (Hussain et al. 2013). Interestingly, Y RNAs were discovered to be modified with N-glycans (Flynn et al. 2019). Substantial research is needed to reveal all the modifications that are likely present in these RNAs.

The presence of mRNAs in the HITS-CLIP-seq dataset was surprising. In recent years, much research has focused on mRNA modifications. The modifications discovered are located in both codong and non-coding mRNA regions. These modifications tend to be simple methylations, although more bulky modifications such as 5-hydroxymethylcytidine or N^6^-acetylcytidine have also been identified (Anreiter et al. 2020; Fu et al. 2014; Arango et al. 2018). The only known uridine modification on mRNA is pseudouridine (Safra et al. 2017). Despite the growing understanding of the variety of mRNA modifications, it seems rather unlikely that ALKBH8 would exhibit its catalytic activity on protein-coding genes. Nevertheless, the mRNAs identified by HITS-CLIP-seq had high read counts and displayed one or two sharp peaks. At the moment, the significance of these data remains enigmatic.

Taken together, our results confirmed ALKBH8 as an enzyme that primarily targets tRNAs that contain a wobble uridine. Among them, we identified tRNA^Lys^(UUU) as a target that had not been previously observed in humans. HITS-CLIP-seq as well as RIP-seq suggest more RNA binding partners of ALKBH8, in particular C/D box snoRNAs. Further studies will reveal the role of these interactions.

## Acknowledgements

We want to thank Karolina Vavrouskova for excellent technical support, Dasa Zigackova for help with cell fractionation, Hana Macickova Cahova for helpful advice on tRNA modifications, and Martin Demko from CEITEC Bioinformatics Core Facility for help with data analysis. This work was supported by the Czech Science Foundation (19-21829S to S.V., T.S. was supported from 20-19617S), by the Ministry of Education, Youth and Sports within programme INTER-COST (LTC18052) and the institutional support CEITEC 2020 (LQ1601). M.B. was supported by Brno City Municipality Scholarship for Talented Ph.D. Students. We further acknowledge the CELLIM core facility supported by MEYS CR (LM2018129 Czech-BioImaging). Computational resources were provided by the project “e-Infrastruktura CZ” (e-INFRA LM2018140) provided within the program Projects of Large Research, Development, and Innovations Infrastructures.

## Data Availability

The high-throughput sequencing datasets were deposited in the NCBI under the number PRJNA678596.

## Supplementary Data

Supplementary data are available for this article.

## Author Contributions

M.B. and S.V. conceived the project. M.B. prepared vectors, stable cell lines and performed CLIP. I.P. performed IF, WB, RIP and northern blot. T.S. performed bioinformatics analyses. P.R. and A.D. purified tRNAs, M.C.S.-D. and M.H. performed LC-MS analyses. I.P. and S.V. interpreted the results and wrote the manuscript. I.P. prepared the figures. S.V. supervised the project.

## Figure Titles and Legends

**FIG. S1.** (A) Chemical structures of the wobble uridine modifications. (B) A schematic view of the domain composition of ALKBH8. RRM is the RNA recognition motif, AlkB-like is the alpha-ketoglutarate-dependent dioxygenase-like domain, and MTase is the S-adenosyl-L-methionine (SAM) dependent methyltransferase domain. (C) Immunoprecipitation of FLAG-ALKBH8 by anti-FLAG antibody during RIP. Protein presence verified by western blot with anti-FLAG antibody. (D) Proportions of CPM normalized values of different non-templated 3 terminal extensions in selected highly represented tRNAs in the RIP experiment. The color code is explained above the graph.

**Fig. S2.** LC-MS analysis uncovered the presence of mcm^5^U and mcm^5^s^2^U in tRNA^Lys^(UUU). (A) Affinity precipitation of tRNA^Glu^(CUC) and tRNA^Lys^(UUU). Denaturing gel analysis of purified tRNAs. RNAs were stained with ethidium bromide. The fragment corresponding to the migration of the purified tRNA was excised from the gel, eluted, and the purified tRNA was subjected to LC-MS analysis. (B) Mass spectrometric analysis at the nucleoside level of the tRNA^Lys^(UUU) isolated from HEK293T cells. The extracted ion chromatograms of mcm^5^s^2^U (left) and mcm^5^U (right) are shown.

**Table S1.** List of primers, linkers, and probes used in this study.

**Table S2.** HITS-CLIP tRNAdb contains CPM-normalized numbers of HITS-CLIP-seq reads mapped by BWA to the tRNA database. HITS-CLIP Human Genome contains CPM-normalized numbers and p-values of HITS-CLIP-seq reads mapped by STAR to the human genome database. Only the reads that remained after mapping to the tRNA database were used for this mapping. RIP-seq tRNAdb and RIP-seq Human Genome are analogous to the previous tables but display data from RIP-seq.

