## Supplementary figures and images for "HITS-CLIP analysis of human ALKBH8 points towards its role in tRNA and noncoding RNA regulation"

### Supplemental Figure 1

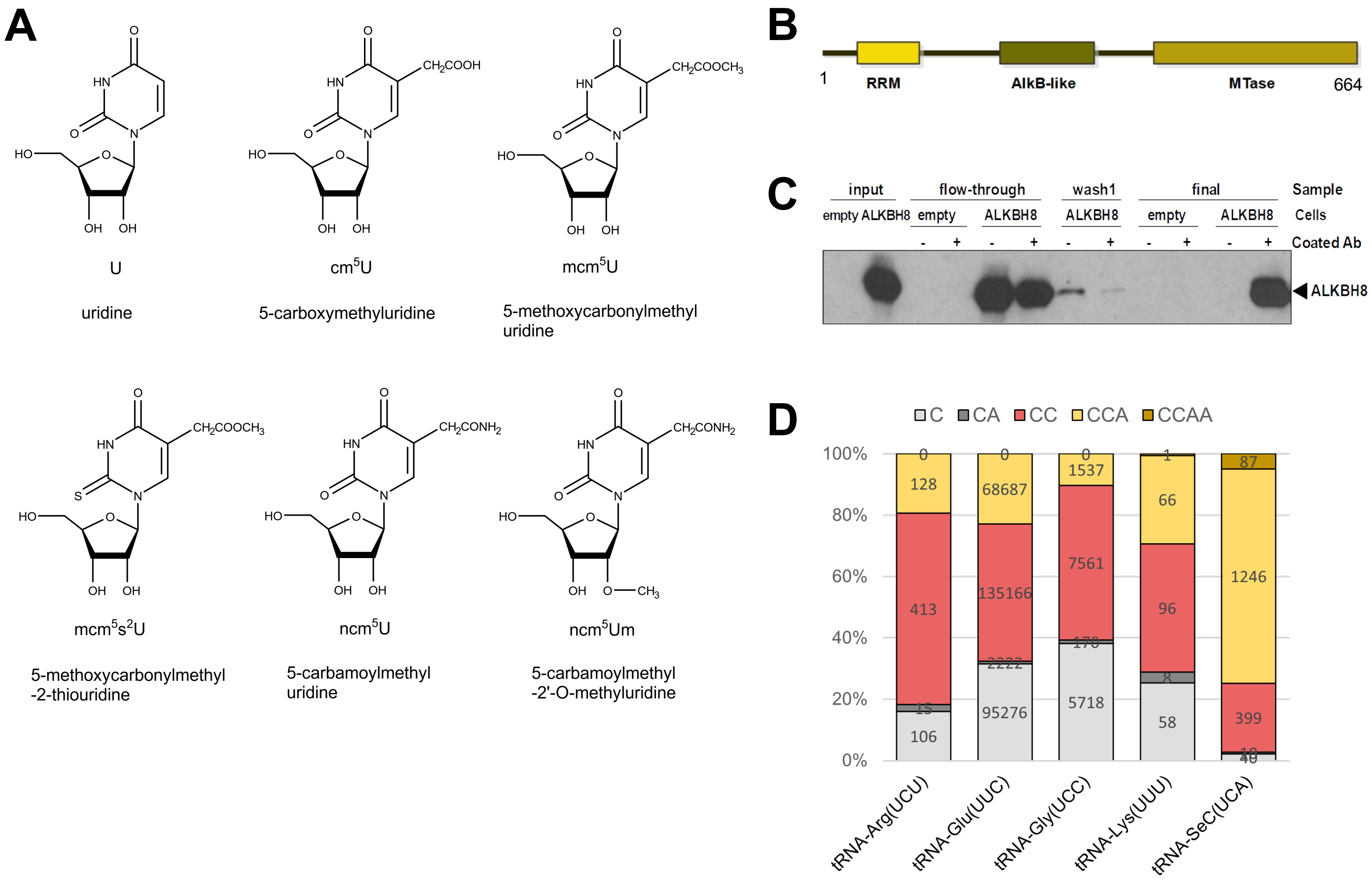

### Supplementary Figure 2

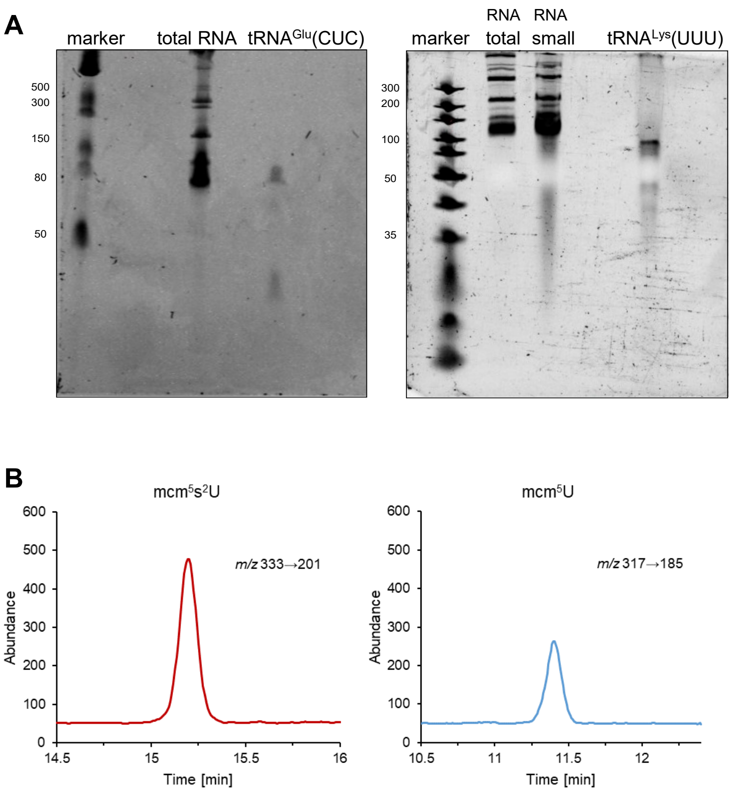
