## Supplementary Table 1 for "HITS-CLIP analysis of human ALKBH8 points towards its role in tRNA and noncoding RNA regulation"

**Table S1**

| Oligonucleotides | Sequence (5' to 3') |
|---|---|
| 3 x FLAG tag | ATGGACTACAAGGACCACGACGGTGACTACAAGGACCACGACATCGACTACAAGGACGACGACGACAAG |
| ALKBH8_fwd | CGGGTACCATGGACAGCAACCATCAAAGTA |
| ALKBH8_rev | CTGGCGGCCGCGGCCTTTTGAAGAATCACAC |
| L3 DNA linker | AppAGATCGGAAGAGCACACGTGTddC |
| L5 RNA linker | OH-AGGGAGGACGAUGCGG-OH |
| TruSeq_adaptor_fwd | AATGATACGGCGACCACCGAGATCTACACTCTTTCCCTACACGACGCTCTTCCGATCTAGAGTTCTACAGTC |
| TruSeq_adaptor_rev | CAAGCAGAAGACGGCATACGAGATCGTTTCACGTGACTGGAGTTCAGACGTGTGCTCTTCCGATC |
| tRNA-Glu-TTC | CCAGGAATCCTAACCGCTAGACCATAT |
| tRNA-Lys-TTT | ACTTGAACCCTGGACCCTCAG |
| tRNA-Lys-CTT | CGCCCAACGTGGGGCTCGAACCCAC |
| tRNA-Gly-CCC | CATGATACCACTACACCAGCGGCGC |
| 5'-tRNA-Glu-CTC_capture | 5AmMC6 CGCCGAATCCTAACCACTAGACCACCA |
| 3'-tRNA-Lys-TTT_capture | 5AmMC6 GCCCGAACAGGGACTTGAACCCTGGAC |
